# Using Hierarchical Similarity To Examine The Genetics of Behçet’s Disease

**DOI:** 10.1101/2021.04.06.438717

**Authors:** Samuel J Shenoi, Erich J Baker

## Abstract

Behçet’s disease (BD) is a multisystem inflammatory disease that affects patients along the historic silk road. Thus far, the pathogenesis of the disease has proved elusive due to the complex genetic interactions and unknown environmental or viral triggering factors of the disease. In this paper, we seek to clarify the genetic factors of the disease while also uncovering other diseases of interest that present with a similar genotype as BD. To do this, we employ a convergent functional genomics approach by leveraging the hierarchical similarity tool available in Geneweaver. Through our analysis, we were able to ascertain 7 BD consensus genes and 16 autoimmune diseases with genetic overlap with BD. The results of our study will inform further research into the pathogenesis of Behçet’s Disease.

## 1 Introduction

Behçet’s disease (BD) is a multi-system inflammatory disease [1]. It was first described by Hulusi Behçet as a disease demonstrating lesions of the mouth, the genitalia, and the eye [2]. The disease is thought to be triggered in genetically pre-disposed individuals by a combination of viral infection and environmental factors, but the exact mechanism of infection is unknown [3].

The genetics of BD are of particular interest. In virtually every studied population, the HLA-B51 gene has been the most closely associated risk factor for BD [4]. However, a sizable portion of BD patients present as HLA-B51 negative, and, conversely, the high prevalence of HLA-B51 in American Indian populations where BD is rare indicates the role of at least one other genetic loci as a factor of BD pathogenesis [4, 5]. Some potential candidates to fill this role have been identified including ERAP1, TNF-*α*, and IL-10 [6, 7, 8, 9, 10, 11, 12, 13]. The lack of a clear mechanism of BD pathogenesis and the genetic variation present in BD patients makes it difficult to separate true genetic factors of the disease from background genetic noise.

The classification of BD as an autoimmune disease and the discovery of isolated shared genetic factors between other autoimmune diseases and BD is intriguing. For instance, ERAP1 has been shown to be a susceptibility gene in BD, psoriasis, and ankylosing spondylitis. Likewise, IL-10 has been associated with multiple other autoimmune diseases, including rheumatoid arthritis, systemic sclerosis, Kawasaki disease, Sjogren’s syndrome, Grave’s disease, myasthenia gravis, psoriasis, autoimmune lymphoproliferative syndrome, and BD [10, 12, 14, 15]. The molecular and genetic overlap between autoimmune diseases presents as a significant area for research, as gaining a deeper understanding of the shared genetic pathways of autoimmune diseases can have important implications for diagnosis, treatments, and future research[16].

Addition research posits an intriguing approach towards solving these problems. In addiction research single genes rarely impact clinical phenotype, and hundreds of variants are needed to fully explain underlying genetics of the diseases. Convergent functional genomics (CFG) is leveraged to isolate important signals against this background. The premise of CFG is straight forward: the more lines of evidence for a gene, the higher it is prioritized as a gene of interest [16, 17]. This approach allows data from sources such as Genome Wide Association Studies (GWAS), gene expression studies, and animal models to be synthesized as evidence to ascertain the impact of genes on phenotype [17]. Even data from sources with limited sample size or lack of replication studies can be used as data points in the CFG approach since it is the collective evidence that together makes up the prioritization list.

To implement the CFG approach, we leverage the computational power of the Geneweaver Hierarchical Similarity (HiSIM) graph [18]. This tool creates a graph that is a hierarchical network of multiway geneset intersections. The resulting geneset interaction graph enables users to find genes connected to all populated subsets of an input set of gene lists (genesets) [19]. The tool takes advantage of a bipartite data structure in order to dynamically enumerate maximal bicliques arranged into hierarchical associations [18, 19]. Nodes at the top of the graph have a limited number of genes but many genesets, while nodes at the bottom of the graph have many genes but fewer genesets as evidence.

In this work, we use the CFG approach to first find candidate genes of interest for BD and evaluate other autoimmune diseases for genetic overlap with BD.

## 2 Methods

### 2.1 Data Collection

Genes associated with BD were collected as genesets on Geneweaver [19]. Each gene-set contained a record of BD associated genes from a single study or source. Sixteen BD genesets were collected overall. Eleven of the genesets originated from GWAS studies and were collected by searching the GWAS Catalog and publicly available curated genesets from Geneweaver [20, 19]. The combined GWAS data came from a global population that included samples from Iranian, Japanese, Turkish, Korean, Spanish, Western European, Middle Eastern, and Han Chinese populations. [21, 22, 23, 24, 25, 26, 27, 28, 29, 24]. One of the GWAS studies collected data regarding BD and a special type of BD that effects the GI tract, Intestinal BD (IBD) [24]. This data source was split into two genesets for purposes of this study. Another geneset was created from the NCBI gene2mesh tool and was included in the study[30]. The Online Mendelian Inheritance in Man (OMIM) database provided an additional geneset tagged “autoinflammatory, familial, Behçet’s-like” [31]. Another geneset was created using information compiled in Malacards [32]. The last two genesets came from a BD review paper that consolidated genes of interest and a BD gene expression profile paper [8, 33]. Finally, we noticed that one of the existing BD genesets on Geneweaver did not include all of the BD associated genes identified by the cited GWAS study. To compensate for this, we created a new geneset from the same GWAS study and added all of the BD associated genes identified by the study to the geneset [34]. The union of all 16 genesets was subsequently collected using the Boolean Algebra tool on Geneweaver and stored as another geneset [19]. Genesets for twenty-seven conditions were compiled in order to test BD’s relation to other autoimmune diseases. Disease genesets were created by using the Geneweaver Boolean Algebra tool on publicly available curated genesets [19]. Due to the large data size, only human genes were used for this run. The list of conditions and number of genes present in the genesets can be seen in Table 1.

**Table 1.** Genetic information from 27 autoimmune diseases were collected from multiple publicly available genesets on Geneweaver. Genesets from each condition were then consolidated into one geneset using the Boolean Algebra tool.

| Condition | Number of Genes | Number of Genesets | Publications |
| --- | --- | --- | --- |
| Achalasia | 12 | 3 | [63] |
| Vitiligo | 107 | 10 | [64, 65, 66, 67, 68, 69, 70] |
| Celiac Disease | 428 | 22 | [71, 72, 73, 74, 75, 76, 77, 78, 63, 79, 80, 81, 82, 83, 84] |
| Crohn's Disease | 273 | 25 | [85, 86, 87, 88, 76, 89, 90, 91, 92, 93, 94, 95, 96, 97, 98, 99, 100, 101, 102] |
| Psoriasis | 382 | 23 | [103, 104, 105, 105, 106, 107, 108, 109, 110, 111, 112, 113, 114, 115, 116, 117] |
| Diabetes | 130 | 20 | [118, 119, 120, 121, 122, 123, 124, 125, 126, 127, 128, 129, 130, 131, 132, 133, 134, 135, 136, 137] |
| Uveitis | 130 | 7 | [63] |
| Sarcoidosis | 107 | 7 | [138, 89, 139] |
| Myositis | 82 | 5 | [63, 140] |
| Scleroderma | 148 | 9 | [141, 142, 63, 143, 144] |
| Agammaglobulinemia | 39 | 3 | [63] |
| Alopecia Areata | 26 | 3 | [145] |
| Restless Leg Syndrome | 27 | 6 | [63, 146, 147, 148, 149] |
| Sjogren Syndrome | 145 | 7 | [150, 128, 151, 152] |
| Hashimoto's Thyroiditis | 99 | 6 | [63, 153, 154] |
| Addison Disease | 40 | 4 | [155] |
| Amyloidosis | 58 | 12 | [63] |
| Ankylosing Spondylitis | 132 | 6 | [156, 157, 79, 158] |
| Chagas Syndrome | 159 | 7 | [159] |
| MS | 201 | 25 | [160, 161, 162, 163, 164, 165, 166, 167, 168, 169, 170, 171, 172, 173, 174, 175, 176, 177, 178, 179, 180, 181, 182, 183] |
| Endometriosis | 226 | 9 | [184, 185, 63, 186, 187, 188, 189] |
| Ulcerative Colitis | 386 | 23 | [190, 191, 192, 193, 194, 195, 89, 196, 92, 197, 198, 199, 200, 97, 201, 202, 203, 204, 205, 206, 207] |
| Hemolytic Anemia | 261 | 20 | [63, 208] |
| Systematic Lupus | 269 | 25 | [209, 210, 211, 212, 150, 213, 214, 215, 216, 217, 218, 219, 220, 221, 222, 223, 143, 224, 225, 226, 227, 228] |
| Liver Cirrhosis | 130 | 8 | [229, 230, 231, 63, 232, 233, 234] |
| Graves' Disease | 121 | 8 | [235, 236, 237, 63, 238, 154, 239] |
| IgA nephropathy | 39 | 6 | [240, 241, 242, 243, 244, 245] |

### 2.2 BD HiSIM Run

To find the consensus genes of BD between all of the collected data sources, the Geneweaver HiSIM graph was run with the 16 BD genesets as input. The parameters used to run the HiSIM graph are reported in Table 2.

**Table 2.**
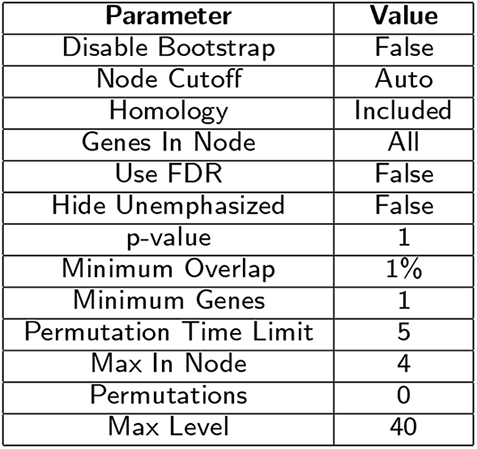
Parameters used for both runs of the Geneweaver HiSIM graph

### 2.3 Autoimmune HiSIM Run

To determine the genetic overlap between BD and other autoimmune diseases. The HiSIM graph tool was run on 27 autoimmune disease genesets constructed using the Boolean Algebra tool and the BD Union geneset. The parameters used to run the HiSIM graph are in Table 2.

The data from the Autoimmune HiSIM run was then filtered to find relevant connections between BD and other autoimmune diseases. The resulting graphs were then plotted using the igraph python package [35] and the BoutrosLab.plotting.general R package [36, 37]

### 2.4 Jaccard Geneset Analysis

After conducting the HiSIM run, a Jaccard Geneset analysis was conducted on autoimmune diseases identified as presenting with a high BD similarity. This analysis determines whether the geneset overlap identified in the Autoimmune Disease Run was statistically significant. The Jaccard Geneset analysis was run using the Jaccard Similarity tool on Geneweaver [19].

### 2.5 Neighbor Joining Analysis

In order to find the top five most genetically similar diseases to BD, a neighbor joining tree was created using the ape R package [38]. The distance between any two diseases was defined using the normalized Jaccard formula seen in Formula 1.

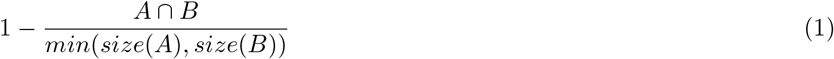

The formula took into account the differences in set size between the two sets in order to determine their overlap. By taking the size of the intersection between the two sets and dividing it by the largest possible overlap that could occur between the two sets, the formula allowed sets of different sizes to be compared. The distance between all possible pair combinations of the 16 genes was calculated and stored in a distance matrix. The distance matrix was then used as input to the neighbor joining program from the ape R package[38, 36].

## 3 Results

### 3.1 BD HiSIM Run

Running the HiSIM graph on the BD collected genesets resulted in the graph shown in Figure 1. Genes found in more than 3 genesets are displayed in Table 3. HLA-B was identified in 7 genesets making it the most common gene amongst all tested genes. IL-10 was the next most common gene and was found in 6 genesets. IL23R was the third most common gene, found in 5 genesets. Finally, HLA-A, STAT4, MICA, and ERAP1 were all found in 4 genesets.

**Table 3.**
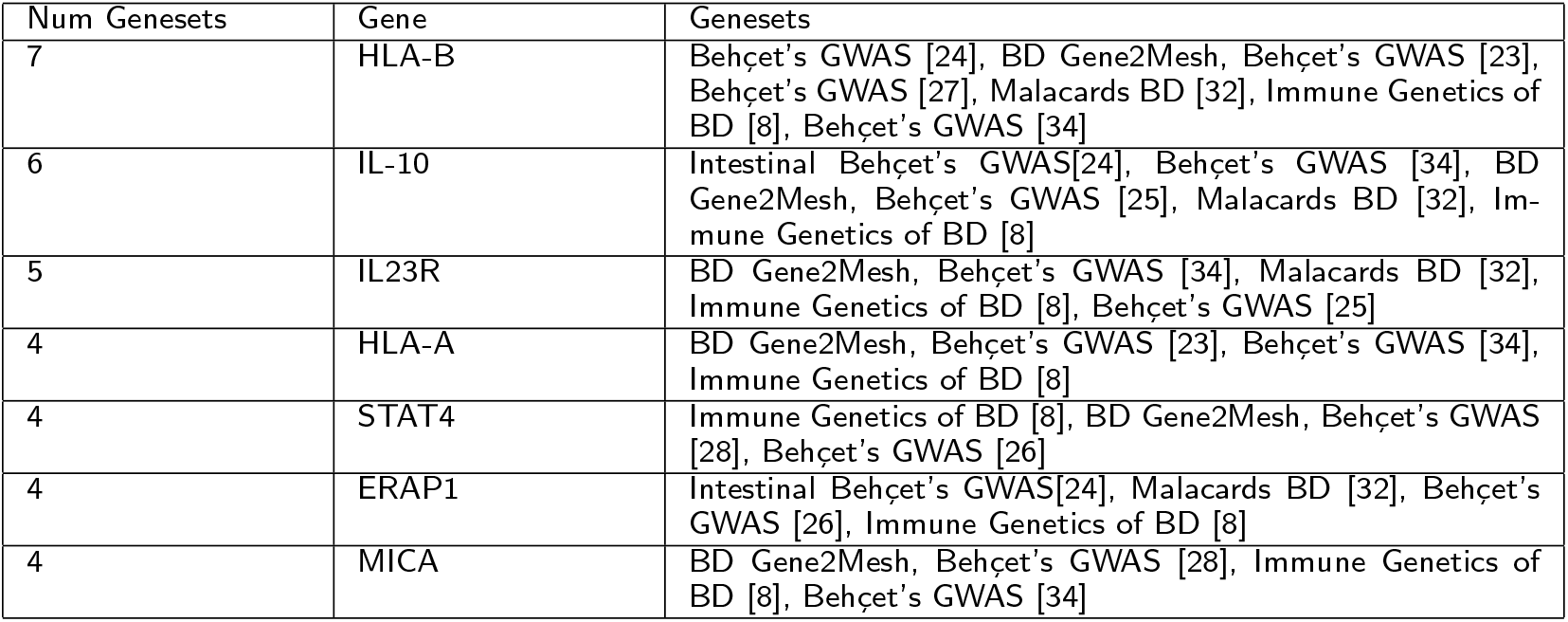
Abbreviated results from the BD HiSIM run. IL10 was identified in 7 genesets making it the most common gene amongst all tested genes. HLA-B was the next most common gene and was found in 6 genesets. IL23R was the third most common gene; it was found in 5 genesets. Finally, HLA-A, STAT4, MICA, and ERAP1 were all found in 4 genesets.

**Figure 1.**
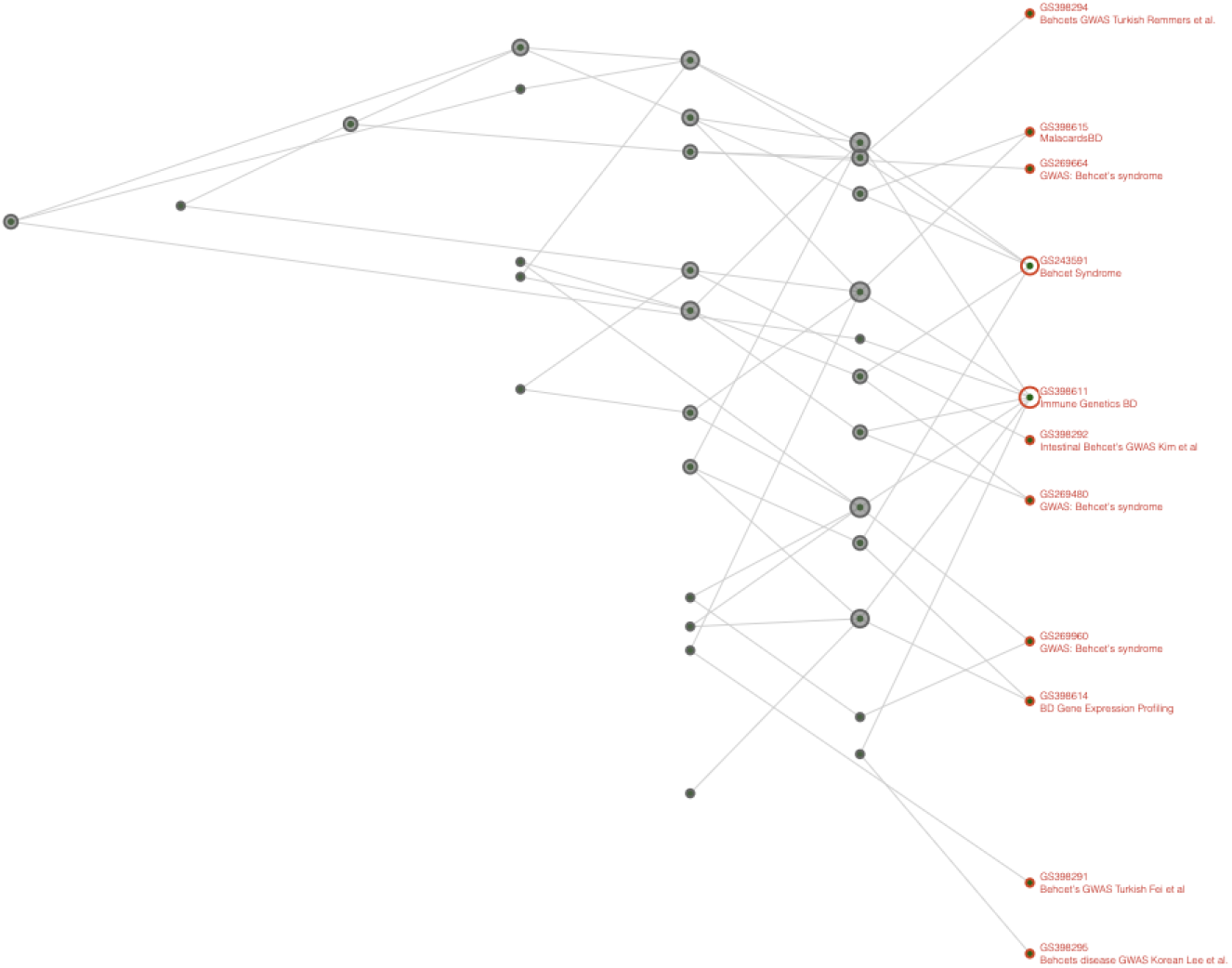
HiSIM Graph between 17 BD gensets.

### 3.2 Autoimmune Disease Run

After collecting autoimmune genesets on Geneweaver, the HiSIM graph was run on 27 unique autoimmune diseases and BD. Figure 2 displays the results of this run. In Figure 2A, all edges between computed nodes are included. The resulting dataset was subsequently filtered in order to remove all nodes that are unrelated to BD; Figure 2B displays the results of filtering.

**Figure 2.**
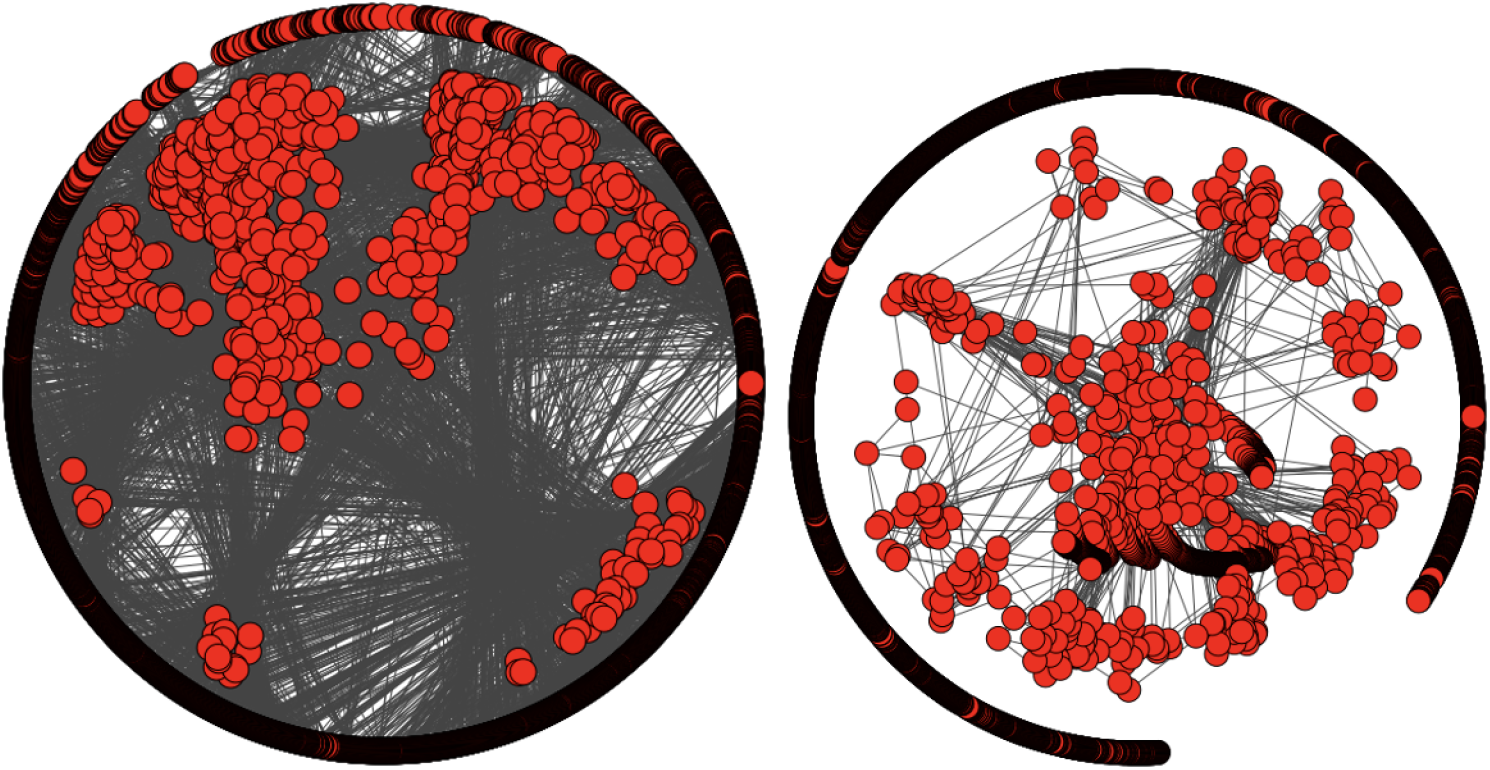
Results from the Autoimmune HiSIM Graph run. 27 autoimmune diseases were compared to BD in order to find which diseases had the largest genetic overlap with BD. **A)** All edges between computed nodes were drawn. **B)** Edges were filtered such that only edges that connected to nodes containing the BD geneset were drawn.

Figure 3 displays the genetic overlap between the autoimmune diseases and BD as determined by the HiSIM graph. Out of the 27 conditions tested, only 16 were found to have some genetic overlap with BD. The 16 identified candidates were then used to subsequently preform the Jaccard Analysis.

**Figure 3.**
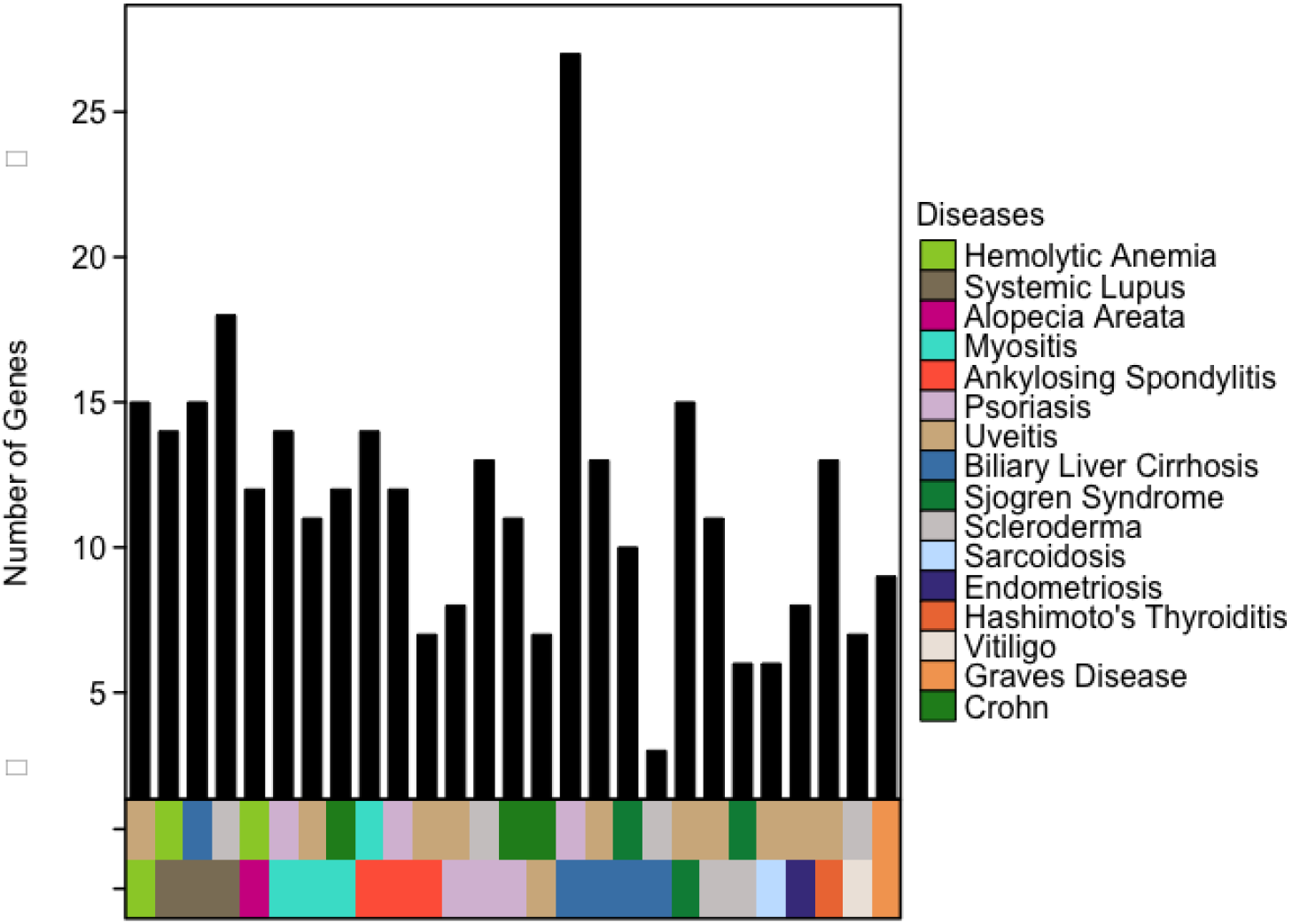
Graph displaying identified HiSIM nodes with the largest gene overlap with BD. 16 conditions were found to have some overlap with BD out of the 27 tested.

### 3.3 Jaccard Geneset Analysis

Using the 16 identified conditions from the Autoimmune HiSIM run, a Jaccard Geneset Analysis was run. The results from the Jaccard Geneset analysis found that the overlap between all of the conditions and BD did not reach a level of statistical significance(Figure 4).

**Figure 4.**
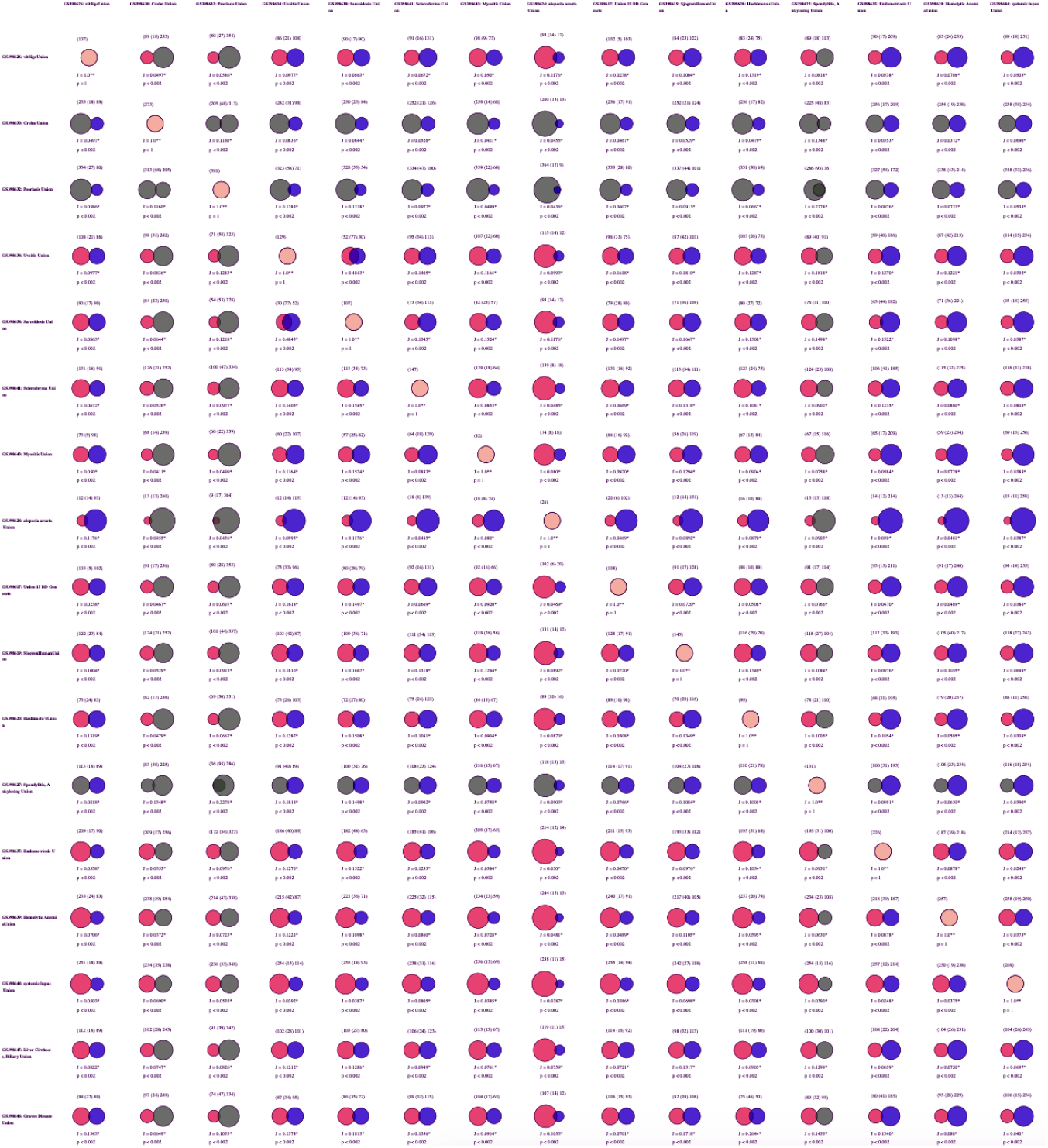
Results of the Jaccard Geneset Analysis. None of the 16 autoimmune disease genesets were identified as having a statistically significant overlap with BD.

### 3.4 Neighbor Joining Analysis

The genetic overlap between BD and the 16 identified autoimmune diseases was then normalized using Formula 1. The normalized genetic overlap was then used as input for the Neighbor Joining Tree, the results of which can be seen in Figure 5. The tree indicates that BD is closest to Sarcoidosis, Uveitis, Sjogren’s Syndrome, Hemolytic Anemia, and Myositis.

**Figure 5.**
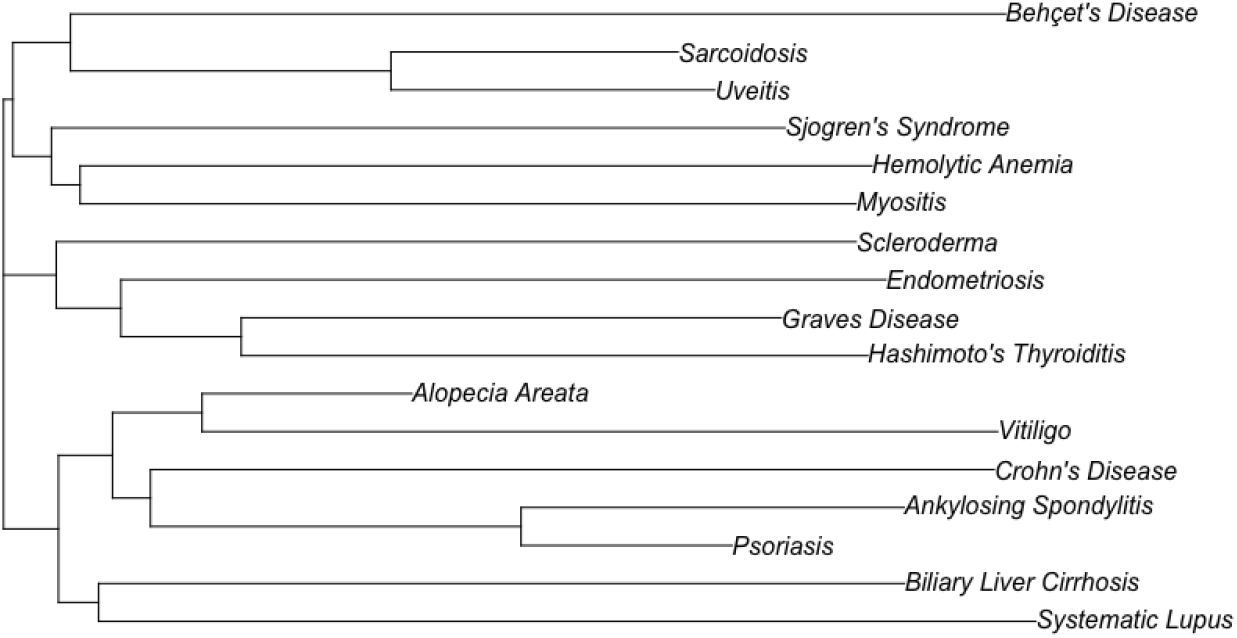
Results of the Neighbor Joining Tree.

## 4 Discussion

The results of the BD HiSIM graph run are consistent with previous BD research. All top scoring genes have previously been identified as having an association with BD [39, 8, 10]. The emergence of HLA-B as the top scoring gene is unsurprising given the history of HLA-B51 as a susceptibility gene of BD and its reputation as the primary gene of interest in BD [4, 6, 39].

The results of the BD autoimmune disease run identified 16 diseases with at least some genetic overlap with BD. While these associations were unable to achieve statistical significance based on the results of our Jaccard Similarity run (Figure 4), we believe that these results can be potentially attributed to the limitations of the Jaccard Similarity test. It has been documented that the Jaccard coefficient, which is used to calculate the Jaccard Similarity, is strongly affected by dataset size [40]. This represents a limitation to our current study that can be addressed in future work.

Additional limitations to our study are primarily centered around data collection. Due to the limited genomics literature available for Behçet’s Disease we were only able to collect 16 unique genesets for analysis. While sufficient for our current study, the addition of future GWAS and other genomic studies will help further elucidate the complex genomic interactions at play in BD. Similarly, our autoimmune disease HiSIM run, while large, is not exhaustive. All of the conditions collected were relatively common and data for these runs were limited to publicly available genesets already present in Geneweaver. This selection criteria excluded non-autoimmune conditions, such as cervical cancer, oral cavity cancer, and oropharyngeal cancer, that might mimic BD phenotype presentations.

To circumvent these limitations, we proposed a modified Jaccard similarity formula (Formula 1) to determine the genetic similarity between any two diseases. We then used this formula to construct a neighbor joining tree which allowed us to identify how genetically similar each of the 16 identified diseases were to BD.

The results of our CFG approach have the benefit of identifying autoimmune diseases with a shared genetic architecture, and is a promising strategy for identifying genes associated with BD. The inclusion of certain diseases in 16 identified diseases was expected. Most notably, Uveitis’s high genetic overlap with BD was expected given that this condition is a symptom of BD [6]. Sjogren’s Syndrome’s inclusion in our list was expected as well given that Sjogren’s Syndrome often presents with other autoimmune diseases [41, 42]. Because of the multiple reports of Sjogren’s Syndrome and BD, Sjogren’s Syndromes association with BD has already been tested and no association between the two was found [43, 42].

Of interest was the close proximity of Sarcoidosis and BD in the neighbor joining tree (Figure 5). Historically, there have been very few reports of patients presenting with both BD and Sarcoidosis [44, 45], but there has been a recent uptick in BD patients presenting with both Sarcoidosis and BD after treatment with TNF antagonists [44]. Clinically, it can be difficult to distinguish between BD and Sarcoidosis considering the similarity of the articular, neurological, and skin lesions between the two diseases [45].

Additionally, the inclusion of psoriasis in our list was noteworthy as well. Psoriatic arthritis has been documented to sometimes be confused with BD articular involvement [46]. Hahn *et al.* recently found that psoriasis patients were twice as likely to be diagnosed with BD based on an analysis of patient data from the National Health Insurance Database of Korea [47]. Finally, Intestinal BD is sometimes misdiagnosed as Crohn’s Disease and vice versa due to the similarity of symptoms, so its inclusion in our list of related diseases was unsurprising[48].

These findings, coupled with our genetic analysis, might provide evidence for a related underlying genetic mechanism related to the pathogenesis of these diseases. It will be left up to future work to fully clarify these relationships.

There have been documented case reports of patients presenting with both BD and one of the other diseases identified from our analysis. There have been a handful of cases of patients presenting with both BD and Sclerosis [49, 50], Myositis [51, 52, 53, 54], Endometriosis [55], Graves’ disease [56], Vitiligo [57, 58], Hemolytic Anemia [59], tuberculous thyroiditis [60, 61],and Biliary Liver Cirrhosis [62]. However, the inclusion of Alopecia Areata in this set of conditions was surprising. To our knowledge, there have been no reported cases of patients presenting with both Alopecia Areata and BD nor evidence of an association between BD and Alopecia Areata.

## 5 Conclusion

Behçet’s disease is a complex multi-system inflammatory disease in which the exact pathogenesis continues to elude researchers [3, 1]. Genetically, the identification of HLA-B51 as a major, but not sole, susceptibility gene has led to the hunt for other genetic factors of the disease [4]. The resulting identification of multiple genes of interest, however, does not explicitly establish the contributions of each of these genes towards the overall presentation of the disease [6, 7, 8, 9, 10, 11, 12, 13]. Furthermore, research into these genes has uncovered roles of these genes in the progression of other autoimmune diseases [10, 12, 14, 15]. It is imperative to research autoimmune diseases in relation to other autoimmune diseases in order to fully understand human disease.

In this study, we employed a functional convergent genomics approach to discover 1) the genetic factors and 2) related autoimmune conditions of Behçet’s Disease. The power of this approach lies in its ability to synthesize information from multiple genomics data sources [16] and in its recognition of shared genetic factors of autoimmune diseases [10, 11, 12]. The ability to quickly and accurately synthesize this information using the Geneweaver HiSIM graph presents a valuable opportunity for further discovery in this line of research [18]. Furthermore, our results using this approach confirmed existing BD research regarding the genetic factors of the disease and identified 16 autoimmune diseases that share an underlying genetic relationship to BD. Almost all of these associations - the only exception being Alopecia Areata - have documented clinical findings linking them with BD, further providing evidence towards our results. It will be left up to further research to fully uncover the complex genetic interactions underlying these diseases as well as the shared genetic mechanisms between them.

## Author’s contributions

S.S carried out the experiments and wrote the manuscript. E.B. supervised the project.

## Acknowledgements

We thank our colleagues from Baylor University who provided insight and expertise that greatly assisted the research, although they may not agree with all of the interpretations/conclusions of this paper.

